# Comparative genomics of *Verticillium dahliae* isolates reveals the *in planta*-secreted effector protein recognized in *V2* tomato plants

**DOI:** 10.1101/2020.06.16.154641

**Authors:** Edgar A. Chavarro-Carrero, Jasper P. Vermeulen, David E. Torres, Toshiyuki Usami, Henk J. Schouten, Yuling Bai, Michael F. Seidl, Bart P.H.J. Thomma

**Author notes:** These authors contributed equally to this work (Edgar A. Chavarro-Carrero, Jasper P. Vermeulen). These authors contributed equally (Michael F. Seidl, Bart P.H.J. Thomma). Corresponding author: B.P.H.J. Thomma.

## Abstract

Plant pathogens secrete effector molecules during host invasion to promote host colonization. However, some of these effectors become recognized by host receptors, encoded by resistance genes, to mount defense response and establish immunity. Recently, a novel resistance was identified in tomato, mediated by the single dominant *V2* locus, to control strains of the soil-borne vascular wilt fungus *Verticillium dahliae* that belong to race 2. We performed comparative genomics between race 2 strains and resistance-breaking race 3 strains to identify the avirulence effector that activates *V2* resistance, termed Av2. We identified 277 kb of race 2-specific sequence comprising only two genes that encode predicted secreted proteins, both of which are expressed by *V. dahliae* during tomato colonization. Subsequent functional analysis based on genetic complementation into race 3 isolates confirmed that one of the two candidates encodes the avirulence effector Av2 that is recognized in *V2* tomato plants. The identification of Av2 will not only be helpful to select tomato cultivars that are resistant to race 2 strains of *V. dahliae*, as the corresponding *V2* resistance gene has not yet been mapped, but also to monitor adaptations in the *V. dahliae* population to deployment of *V2*-containing tomato cultivars in agriculture.

## INTRODUCTION

In nature, plants are continuously threatened by potential plant pathogens. However, most plants are resistant to most potential plant pathogens due to an efficient immune system that becomes activated by invasion patterns of diverse nature (Cook, et al., 2015; Dangl & Jones, 2001). Throughout time, different conceptual frameworks have been put forward to describe the molecular basis of plant–pathogen interactions and the mechanistic underpinning of plant immunity. Initially, Harold Flor introduced the gene-for-gene model in which a single dominant host gene, termed a resistance (*R*) gene, induces resistance in response to a pathogen expressing a single dominant avirulence (*Avr*) gene (Flor, 1942). Isolates of the pathogen that do not express the allele of the *Avr* gene that is recognized escape recognition and are assigned to a resistance-breaking race. In parallel to these race-specific Avrs, non-race-specific elicitors were described as conserved microbial molecules that are often recognized by multiple plant species (Darvill & Albersheim, 1984). The recognition by plants of Avrs and of non-race-specific elicitors, presently known as microbe-associated molecular patterns (MAMPs), was combined in the ‘zig-zag’ model (Jones & Dangl, 2006). In this model, MAMPs are perceived by cell surface-localized pattern recognition receptors (PRRs) to trigger MAMP-triggered immunity (MTI), while effectors are recognized by cytoplasmic receptors that are known as resistance (R) proteins to activate effector-triggered immunity (ETI) (Jones & Dangl, 2006). Importantly, the model recognizes that Avrs function to suppress host immune responses in the first place, implying that these molecules, besides being avirulence determinants, act as virulence factors through their function as effector molecules (Jones & Dangl, 2006). A more recent model, termed the Invasion Model, recognizes that the functional separation of MTI and ETI is problematic and proposes that the corresponding receptors, collectively termed invasion pattern receptors (IPRs), detect either externally encoded or modified-self ligands that indicate invasion, termed invasion patterns (IPs), to mount an effective immune response (Cook et al., 2015; Thomma et al., 2011). However, it is generally appreciated that microbial pathogens secrete dozens to hundreds of effectors to contribute to disease establishment, only some of which are recognized as Avrs (Rovenich et al., 2014).

Plant invasion pattern receptors (IPR) encompass typical *R* genes which have been exploited for almost a century to confer resistance against plant pathogens upon introgression from sexually compatible wild relatives into elite cultivars (Dangl et al., 2013; Dodds & Rathjen, 2010). Most *R* genes encode members of a highly polymorphic superfamily of intracellular nucleotide-binding leucine-rich repeat (NLR) receptors, while others encode cell surface receptors (Dangl et al., 2013). Unfortunately, most *R* genes used in commercial crops are short-lived because the resistance that they provide is rapidly broken by pathogen populations as their deployment in monoculture-based cropping systems selects for pathogen variants that overcome immunity (Dangl et al., 2013; Stukenbrock & McDonald, 2008). Such breaking of resistance occurs upon purging of the *Avr* gene, sequence diversification, or by employment of novel effectors that subvert the host immune response (Cook et al., 2015; Stergiopoulos et al., 2007).

The molecular cloning of the first bacterial *Avr* gene from *Pseudomonas syringae* pv. *glycinea* was reported in 1984 (Staskawicz et al., 1984), the first fungal *Avr* gene from *Cladosporium fulvum* in 1991 (van Kan et al., 1991) and the first oomycete *Avr* gene from *Phytophthora sojae* in 2004 (Shan et al., 2004). Dozens of additional *Avr* genes have been cloned since then. Initially, such *Avr* genes were identified by reverse genetics and map-based cloning strategies, but also other approaches have been exploited such as functional screens (Luderer et al., 2002; Takken et al., 2000). More recently, advances in (the affordability of) genome sequencing have allowed the cloning of novel *Avrs* by combining comparative genomics or transcriptomics with functional assays, a trend that was spearheaded by the cloning of the first *Avr* gene from *Verticillium dahliae* in 2012 (de Jonge et al., 2012; Mesarich et al., 2014; Schmidt et al., 2016).

*Verticillium dahliae* is a soil-borne fungal pathogen and causal agent of Verticillium wilt on a broad range of host plants that comprises hundreds of dicotyledonous plant species, including numerous crops such as tomato, potato, lettuce, olive, and cotton (Fradin & Thomma, 2006; Klosterman et al., 2009). The first source of genetic resistance toward Verticillium wilt was identified in tomato (*Solanum lycopersicum*) in the early 1930s in an accession called Peru Wild (Schaible et al., 1951). The resistance is governed by a single dominant locus, designated *Ve* (Diwan et al., 1999), comprising two genes that encode cell surface receptors of which one, *Ve1*, acts as a genuine resistance gene (Fradin & Thomma, 2006). Shortly after its deployment in the 1950s, resistance-breaking strains have appeared that were assigned to race 2 whereas strains that are contained by *Ve1* belong to race 1 (Alexander, 1962). Thus, *Ve1* is characterized as a race-specific *R* gene, and resistance-breaking strains have become increasingly problematic over time (Alexander, 1962; Dobinson et al., 1996). With comparative population genomics of race 1 and race 2 strains, the *V. dahliae* avirulence effector that is recognized by tomato Ve1 was identified as VdAve1, an effector that is secreted during host colonization (de Jonge et al., 2012). As anticipated, it was demonstrated that VdAve1 acts as a virulence factor on tomato plants that lack the *Ve1* gene and that, consequently, cannot recognize VdAve1 (de Jonge et al., 2012). Whereas all race 1 strains carry an identical copy of *VdAve1*, all race 2 strains analysed to date are characterized by complete loss of the *VdAve1* locus (de Jonge et al., 2012; Faino et al., 2016). Intriguingly, phylogenetic analysis has revealed that *VdAve1* was horizontally acquired by *V. dahliae* from plants (de Jonge et al., 2012; Shi-Kunne et al., 2019), after which the effector gene was lost multiple times independently, presumably due to selection pressure exerted by the *Ve1* locus that has been introgressed in most tomato cultivars (Faino et al., 2016). Despite significant efforts, attempts to identify genetic sources for race 2 resistance in tomato have remained unsuccessful for a long time (Baergen et al., 1993). Recently, however, a source of race 2 resistance was identified in the wild tomato species *Solanum neorickii* (Usami et al., 2017). This genetic material was used to develop the rootstock tomato cultivars Aibou, Ganbarune-Karis and Back Attack by Japanese breeding companies, in which resistance is controlled by a single dominant locus, denoted *V2* (Usami et al., 2017). However, experimental trials using race 2-resistant rootstocks revealed resistance-breaking *V. dahliae* strains that, consequently, are assigned to race 3 (Usami et al., 2017). In this study, we performed comparative genomics based on Oxford Nanopore sequencing technology, combined with functional assays, to identify the avirulence effector Av2 that activates race-specific resistance in tomato lines that carry *V2*.

## MATERIALS AND METHODS

### *V. dahliae* inoculation and phenotyping

Plants were grown in potting soil (Potgrond 4, Horticoop, Katwijk, the Netherlands) under controlled greenhouse conditions (Unifarm, Wageningen, the Netherlands) with day/night temperature of 24/18°C for 16-h/8-h periods, respectively, and relativity humidity between 50% and 85%. For *V. dahliae* inoculation, 10-day-old seedlings were root-dipped for 10 min as previously described (Fradin et al., 2009). Disease symptoms were scored at 21 days post inoculation (dpi) by measuring the canopy area to calculate stunting as follows:

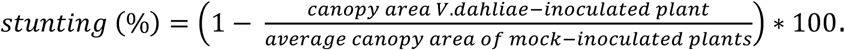

### High Molecular Weight (HMW) DNA isolation and Nanopore sequencing

Conidiospores were harvested from potato dextrose agar (PDA) plates, transferred to Czapek dox medium and grown for ten days. Subsequently, fungal material was collected on Miracloth, freeze-dried overnight and ground to powder with mortar and pestle of which 300 mg was incubated for one hour at 65°C with 350 µL DNA extraction buffer (0.35 M Sorbitol, 0.1 M Tris-base, 5 mM EDTA pH 7.5), 350 µL nucleic lysis buffer (0.2 M Tris, 0.05 M EDTA, 2 M NaCl, 2% CTAB) and 162.5 µL Sarkosyl (10% w/v) with 1% β-mercaptoethanol. Next, 400 µL of phenol/chloroform/isoamyl alcohol (25:24:1) was added, shaken and incubated at room temperature (RT) for 5 minutes before centrifugation at 16,000 g for 15 min. After transfer of the aqueous phase to a new tube, 10 µl of RNAase (10 mg/µL) was added and incubated at 37°C for one hour. Subsequently, half a volume of chloroform was added, shaken and centrifuged at 16,000 g for 5 min at RT, after which the chloroform extraction was repeated. Next, the aqueous phase was mixed with 10 volumes of 100% ice-cold ethanol, incubated for 30 min at RT, and the DNA was fished out using a glass hook, transferred to a new tube, and washed twice with 500 µL 70% ethanol. Finally, the DNA was air-dried, resuspended in nuclease-free water and incubated at 4°C for two days. The DNA quality, size and quantity were assessed by Nanodrop, gel electrophoresis and Qubit analyses.

Library preparation with the Rapid Sequencing Kit (SQK-RAD004) was performed according to the manufacturer’s instructions (Oxford Nanopore Technologies, Oxford, UK) with 400 ng HMW DNA. An R9.4.1 flow cell (Oxford Nanopore Technologies, Oxford, UK) was loaded and run for 24 hours. Base calling was performed using Guppy (version 3.1.5; Oxford Nanopore Technologies, Oxford, UK) with the high-accuracy base-calling algorithm. Adapter sequences were removed using Porechop (version 0.2.4 with default settings; Wick, 2018). Finally, the reads were self-corrected, trimmed and assembled using Canu (Version 1.8; Koren et al., 2017).

### Comparative genomics and candidate identification

Self-corrected reads from *V. dahliae* race 3 strains were mapped against the reference genome using BWA-MEM (version 0.7.17; default settings; Li, 2013). Reads with low mapping quality (score <10) were removed using Samtools view (version 1.9; setting: -q 10) (Li et al., 2009), and reads mapping in regions with low coverage (<10x) were discarded using Bedtools coverage (version 2.25.0; setting: -d) (Quinlan & Hall, 2010). Self-corrected race 2 strain reads were mapped against the retained reference genome-specific regions that are absent from the race 3 strains. Retained sequences shared by the reference and every race 2 strain, while absent from every race 3 strain, were retained as *Av2* candidate regions.

The previously determined annotation of *V. dahliae* strain JR2 (Faino et al., 2015) was used to extract genes when JR2 or TO22 were used as alignment references. To this end, retained sequences shared by the TO22 reference assembly and race 2 strains, absent from race 3 strains, were mapped against the JR2 genome assembly, and genes in the shared sequences were extracted. The remaining sequences that did not map to the *V. dahliae* strain JR2 genome assembly were annotated using Augustus (version 2.1.5; default settings; Stanke et al., 2006). SignalP software (version 4.0; Petersen et al. 2011) was used to identify N-terminal signal peptides.

### Real-time PCR

To determine expression profiles of *Av2* candidate genes during *V. dahliae* infection of tomato, two-week-old tomato (cv. Moneymaker) seedlings were inoculated with *V. dahliae* strain JR2 or TO22, and stems were harvested up to 14 days post inoculation (dpi). Furthermore, conidiospores were harvested from five-day-old PDA plates. Total RNA extraction and cDNA synthesis were performed as previously described (Santhanam et al., 2013). Real time-PCR was performed with primers listed in Table 3, using the *V. dahliae* glyceraldehyde-3-phosphate dehydrogenase gene (*GAPDH*) as endogenous control.

### Genome mining

In total, 44 previously sequenced *V. dahliae* strains and eight strains sequenced in this study were mined for *Av2* gene candidates using BLASTn. Gene sequences were extracted using Bedtools (setting: getfasta) (Quinlan & Hall, 2010) and aligned to determine allelic variation using Espript (version 3.0; default settings) (Robert & Gouet, 2014). Similarly, amino acid sequences were aligned using Espript (Robert & Gouet, 2014).

To determine the genomic localization of *XLOC_00170* and *Evm_344*, the *V. dahliae* strain JR2 assembly and annotation were used (Faino et al., 2015) together with coverage plots from reads of race 3 and race 2 strains as described in comparative genomics approach IV (Table 2) using R scripts, with the package karyoploteR for R (version 3.6) using kpPlotBAMCoverage function. The schematic representation of the genomic region on chromosome 4 with *XLOC_00170* and *Evm_344* was generated using Integrative Genomics Viewer (IGV) software v2.6.3 (J. T. Robinson et al., 2011) and R package (version 3.6) Gviz (Hahne and Ivanek, 2016).

### Phylogenetic tree construction

The phylogenetic tree of 52 *V. dahliae* strains was generated with Realphy (version 1.12) (Bertels et al., 2014) using Bowtie2 (version 2.2.6) (Langmead & Salzberg, 2012) to map genomic reads against the *V. dahliae* strain JR2 assembly. A maximum likelihood phylogenetic tree was inferred using RAxML (version 8.2.8) (Stamatakis, 2014).

### Presence-absence variation analysis

Presence-absence variation (PAV) was identified by using whole-genome alignments for 17 *V. dahliae* strains. Paired-end short reads were mapped to *V. dahliae* strain JR2 (Faino et al., 2015) using BWA-mem with default settings (Li and Durbin, 2009). Long-reads were mapped using minimap2 with default settings (Li, 2018). Using the Picard toolkit (http://broadinstitute.github.io/picard/), library artifacts were marked and removed with -*MarkDuplicates* followed by *-SortSam* to sort the reads. Raw read coverage was averaged per 100 bp non-overlapping windows using the BEDtools *-multicov* function (Quinlan and Hall, 2010). Next, we transformed the raw read coverage values to a binary matrix by applying a cut-off of 10 reads for short-read data; >=10 reads indicate presence (1) and <10 reads indicate absence (0) of the respective genomic region. For long-read data a cut-off of 1 read was used; >=1 read indicates presence (1) and <1 read indicates absence (0). The total number of PAV counts for each of the 100 bp genomic windows within 100 kb upstream and downstream of the candidate effectors was summarized.

### Genetic complementation and functional analysis

For genomic complementation of race 3 strains GF-CB5 and HOMCF, a genomic construct was generated comprising the coding sequence of *XLOC_00170* or *Evm_344* in vector pFBT005 behind the *VdAve1* promoter, using primers CO-XLOC00170-F and CO-XLOC00170-R for *XLOC_00170* or CO-Evm344-F and CO-Evm344-R for *Evm_344* (Table 3).

*A. tumefaciens*-mediated transformation (ATMT) was performed as described previously (Ökmen et al., 2013) with few modifications. *A. tumefaciens* was grown in 5 mL minimal medium (MM) supplemented with 50 µg/mL kanamycin at 28°C for two days. After subsequent centrifugation at 3,000 g (5 min), cells were resuspended in 5 mL induction medium (IM) supplemented with 50 µg/mL kanamycin, adjusted to OD_600_ 0.15 and grown at 28°C for minimum 6 h until OD_600_ 0.5. Simultaneously, conidiospores of *V. dahliae* race 3 strains GF-CB5 and HOMCF were harvested after one week of cultivation on PDA plates with water, rinsed, and adjusted to a final concentration of 10^6^ conidiospores/mL. The *A. tumefaciens* suspension was mixed with *V. dahliae* conidiospores in a 1:1 volume ratio and 200 µL of the mixture was spread onto PVDF membranes in the centre of IM agar plates. After two days at 22°C, membranes were transferred to fresh PDA plates supplemented with 20 µg/mL nourseothricin and 200 µM cefotaxime and incubated at 22°C for two weeks until *V. dahliae* colonies emerged. Transformants that appeared were transferred to fresh PDA supplemented with 20 µg/ml nourseothricin and 200 µM cefotaxime. Successful transformation was verified by PCR and DNA sequencing.

*V. dahliae* inoculations were performed as described previously (Fradin et al., 2009). Disease symptoms were scored ten days after inoculation by measuring the canopy area to calculate stunting when compared with mock-inoculated plants. For biomass quantification, stems were freeze-dried and ground to powder, of which ∼100 mg was used for DNA isolation. Real-time PCR was conducted with primers SlRUB-Fw and SlRUB-Rv for tomato *RuBisCo* and primers ITS1-F and STVe1-R for *V. dahliae* ITS (Table 3). Real-time PCR conditions were as follows: an initial 95°C denaturation step for 10 min followed by denaturation for 15 s at 95°C and annealing for 30 s at 60°C, and extension at 72°C for 40 cycles.

## RESULTS

### Identification of *Verticillium dahliae* strains that escape *V2* resistance

To identify Av2 as the *V. dahliae* gene that mediates <u>a</u>virulence on tomato <u>V2</u> plants, we pursued a comparative genomics strategy by searching for race 2-specific regions that are absent from race 3 strains. To this end, we performed pathogenicity assays with a collection of *V. dahliae* strains on a differential set of tomato genotypes, comprising (I) Moneymaker plants that lack *V. dahliae* resistance genes, (II) *Ve1*-transgenic Moneymaker plants that are resistant against race 1 and not against race 2 strains (Fradin et al., 2009), and (III) Aibou plants that carry *Ve1* and *V2* and are therefore resistant against race 1 as well as race 2 strains (Usami et al., 2017) (Fig 1A). First, we aimed to confirm the race assignment of eight *V. dahliae* strains that were previously tested by Usami et al. (2017) (Table 1). Additionally, three strains that were previously assigned to race 2 were included (de Jonge et al., 2012). Finally, *V. dahliae* strain JR2 was included because of its gapless telomere-to-telomere assembly (Faino et al., 2015).

**Table 1.** *V. dahliae* strains used in this study for comparative genomics and genome assembly statistics.

| Strain | Previous<br>race | Ref.<sup>1</sup> | Novel<br>race | Platform<sup>2</sup> | Data<br>(Gb)<sup>3</sup> | Assembly<br>size (Mb) | No. of<br>contigs<sup>4</sup> | Contig<br>N50 (Kb) | Used in<br>approaches | Ref.<sup>1</sup> |
| --- | --- | --- | --- | --- | --- | --- | --- | --- | --- | --- |
| JR2 | 1 | A | 1 | PacBio | 8.9 | 36.1 | 8 | 4168 | IV | D |
| DVDS26 | 2 | B | 2 | Illumina | 1 | 35.3 | 5,361 | 47.1* | II, III, IV | B |
| DVD161 | 2 | B | 2, 3 | Illumina | 1 | 34.1 | 4,078 | 42.4* | II, III, IV | B |
| DVD3 | 2 | B | 2, 3 | Illumina | 1 | 34.1 | 9,318 | 43.9* | II, III, IV | B |
| TO22 | 2 | C | 2 | Nanopore | 4 | 34.9 | 20 | 12.4 | I, II, III, IV | TS |
| UD1-4-1 | 2 | C | 2 | Nanopore | 1.9 | 34.6 | 18 | 18.1 | I, II, III, IV | TS |
| GF1207 | 2 | C | 2 | Nanopore | 1.6 | 34.8 | 69 | 8.5 | III | TS |
| GF-CA2 | 2 | C | 2 | Nanopore | 2 | 35 | 38 | 9.1 | I, II, III, IV | TS |
| GF-CB5 | 3 | C | 2, 3 | Nanopore | 4 | 34.8 | 19 | 11.4 | I, II, III, IV | TS |
| VT-2A | 3 | C | 2, 3 | Nanopore | 1.8 | 34.8 | 22 | 10.2 | III | TS |
| GF1192 | 3 | C | 2, 3 | Nanopore | 2 | 34.6 | 23 | 14.5 | I, II, III, IV | TS |
| HOMCF | 3 | C | 2, 3 | Nanopore | 2 | 36.1 | 33 | 10.1 | I, II, III, IV | TS |

<sup>1</sup>References: A: Fradin et al., 2009; B: de Jonge et al., 2012; C: Usami et al., 2017; D: Faino et al., 2015; TS: this study.
<sup>2</sup>Sequencing platform used.
<sup>3</sup>Amount of sequencing data generated
<sup>4</sup>For strains DVDS26, DVD161 and DVD3 previously determined scaffold statistics are shown (de Jonge et al., 2012)

**Table 2.** Comparative genomics for four different groupings of race 2 and race 3 strains to reveal race 2-specific *Avr* candidates.

| Approach | I | II | III | IV |
| --- | --- | --- | --- | --- |
| Reference strain | TO22 | TO22 | TO22 | JR2 |
| Race 2 | GF-CA2<br>TO22<br>UD-1-4-1 | GF-CA2<br>TO22<br>UD-1-4-1<br>DVDS26 | GF-CA2<br>TO22<br>UD-1-4-1<br>DVDS26<br>GF1207 | GF-CA2<br>TO22<br>UD-1-4-1<br>DVDS26 |
| Race 3 | GF-1192<br>GF-CB5<br>HOMCF | GF-1192<br>GF-CB5<br>HOMCF<br>DVD161<br>DVD3 | GF-1192<br>GF-CB5<br>HOMCF<br>DVD161<br>DVD3<br>VT-2A | GF-1192<br>GF-CB5<br>HOMCF<br>DVD161<br>DVD3 |
| Retained (kb) | 670 | 660 | 563 | 277 |
| Shared with JR2 (kb) | 282 | 277 | 222 | 277 |
| #JR2 genes | 44 | 41 | 40 | 41 |
| #Augustus genes | 78 | 74 | 70 | -- |
| #Secreted | 8 | 6 | 6 | 2 |
| Retained candidates | • XLOC_00170<br>• evm.model.co ntig1569.344<br>• tig00000058: 1027588-1028906<br>• tig00000058: 1116044-1116494<br>• tig00000151: 403362-404089<br>• tig00017428: 835657-837290<br>• tig00000058: 1220542-1226128<br>• tig00000198: 12352-12816 | • XLOC_00170<br>• evm.model.co ntig1569.344<br>• tig00000058: 1027588-1028906<br>• tig00000058: 1116044-1116494<br>• tig00000151: 403362-404089<br>• tig00017428: 835657-837290 | • XLOC_00170<br>• evm.model.co ntig1569.344<br>• tig00000058: 1027588-1028906<br>• tig00000058: 1116044-1116494<br>• tig00000151: 403362-404089<br>• tig00017428: 835657-837290 | • XLOC_00170<br>• evm.model.co ntig1569.344 |

**Table 3.**
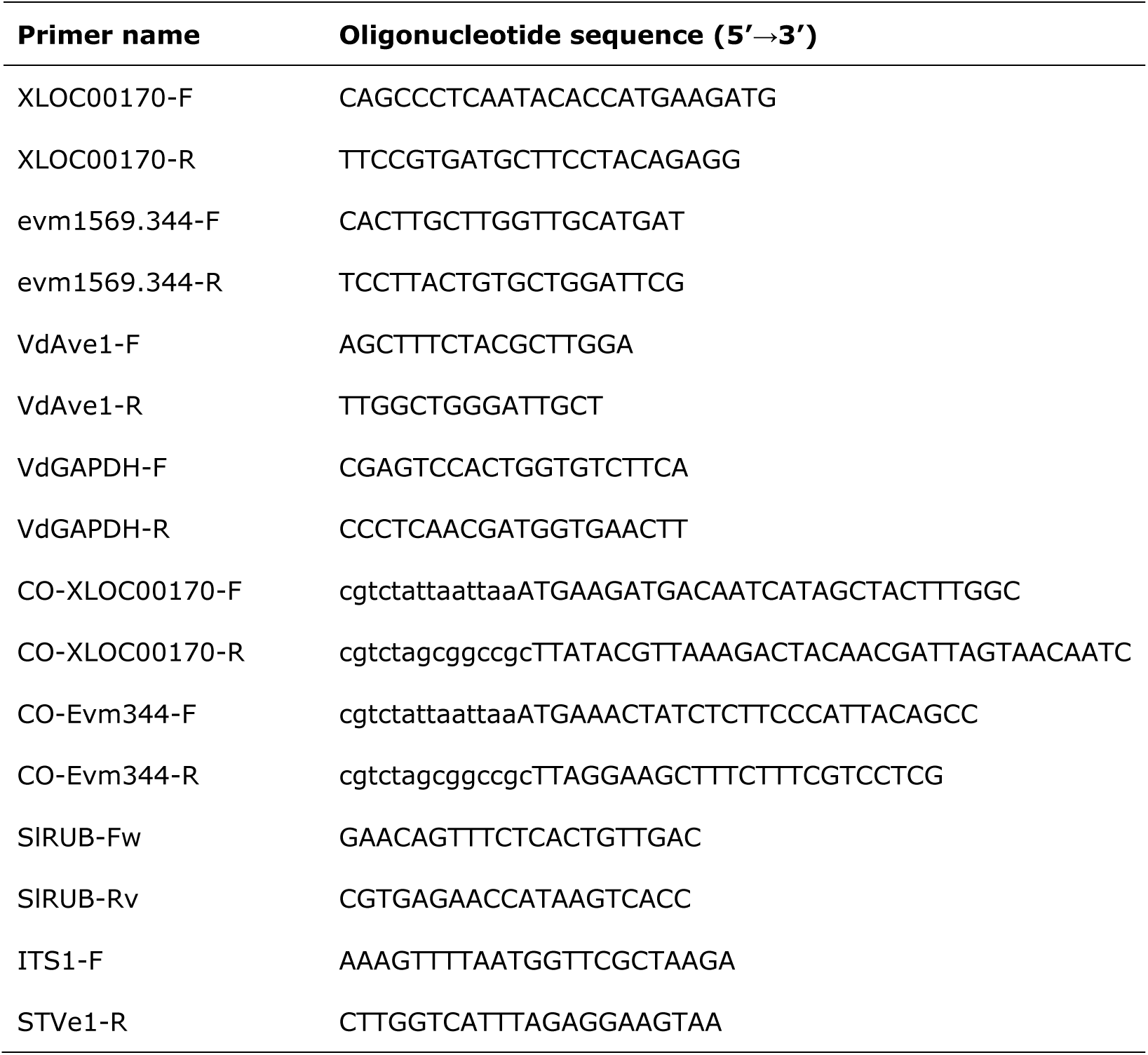
Primers used in this study.

**Figure 1.**
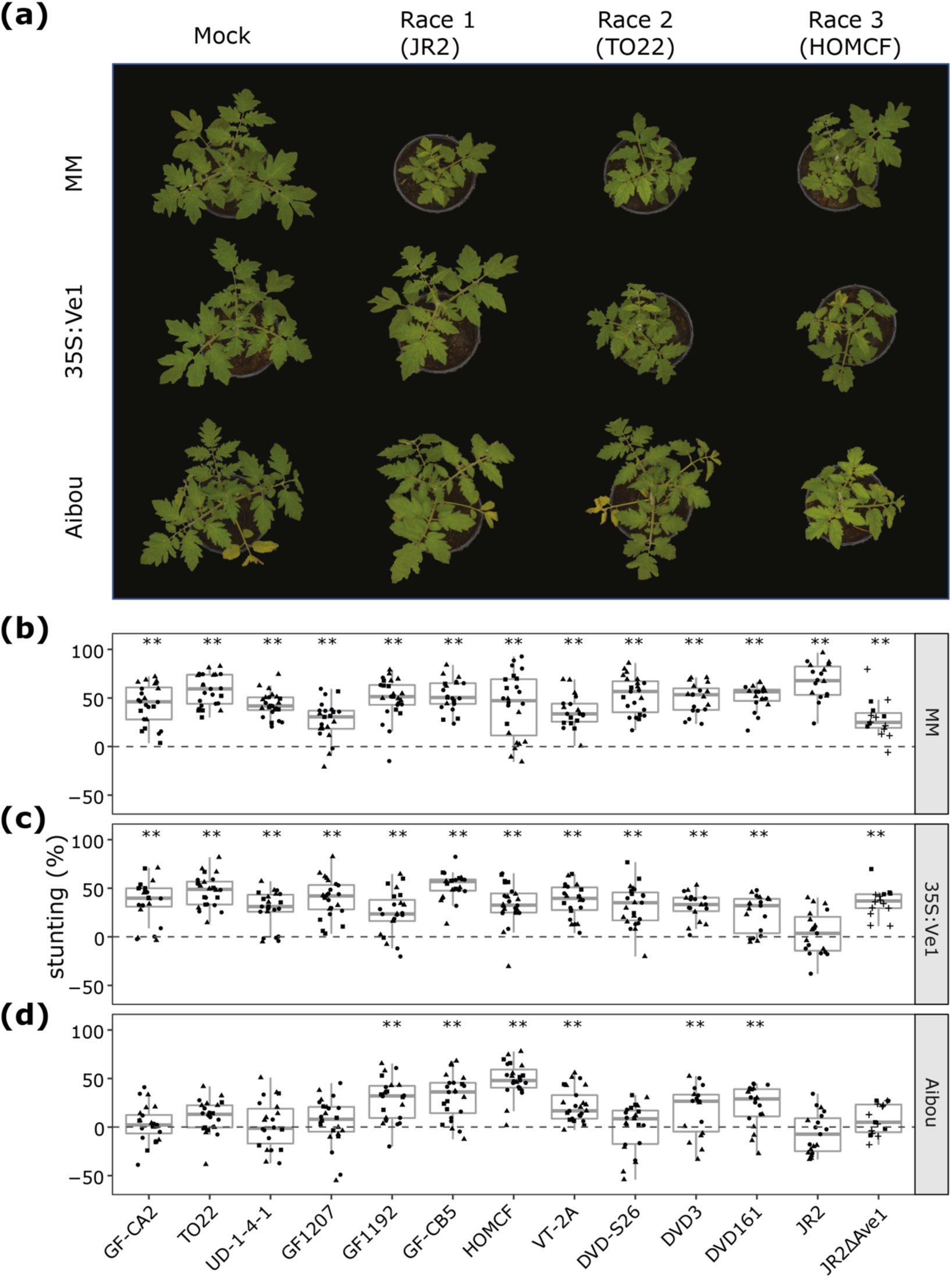
Pathogenicity phenotyping of a collection of *Verticillium dahliae* strains on tomato. **(a)** Typical appearance of *V dahliae* infection by strain JR2, TO22 and HOMCF as representatives for race 1, 2 and 3, respectively on Moneymaker plants that lack known *V. dahliae* resistance genes, *Ve1*-transgenic Moneymaker plants that are resistant against race 1 and not against race 2 or 3 strains, and Aibou plants that carry *Ve1* and *V2* and are therefore resistant against race 1 as well as race 2 strains, but not against race 3 strains at 21 days post inoculation (dpi). **(b-d)** Measurement of *V. dahliae*-induced stunting on wild-type Moneymaker plants **(b)**, *Ve1*-transgenic Moneymaker plants (35S:Ve1) **(c)** and Aibou plants **(d)** at 21 dpi. The graphs show collective data from four different experiments indicated with different symbols (circles, squares, triangles and plus symbols), and asterisks indicate significant differences as determined with an ANOVA followed by a Fishers’ LSD test (p < 0.01).

At three weeks post inoculation, all strains caused significant stunting on the universally susceptible Moneymaker control (Fig. 1a, b), confirming that all strains are pathogenic on tomato. Similarly, except for the race 1 strain JR2, all strains caused significant stunting on *Ve1*-transgenic Moneymaker plants (Fig. 1a, c), corroborating that they lack *VdAve1*. This observation implies that, except for strain JR2, none of the strains is of race 1 and that a potential containment on Aibou plants cannot be caused by Ve1 recognition of the VdAve1 effector. Importantly, most of the strains that were used by Usami and colleagues (2017), and that were previously assigned to race 2, did not cause significant stunting on Aibou, whereas most of the strains that were assigned to race 3 caused clear symptoms of Verticillium wilt disease (Fig. 1a, d). More specifically, this concerned the race 2 strains TO22, UD-1-4-1 and GF-CA2 and the race 3 strains HOMCF, GF-CB5 and GF1192 (Fig. 1; Usami et al., 2017). The exceptions are strains GF1207 and VT-2A that were previously assigned to race 2 and 3, respectively (Usami et al., 2017), but for which the phenotyping in our assays was ambiguous due to a relatively low degree of virulence on Moneymaker plants (Fig. 1b, d). The previously assigned race 2 strain DVDS26 (de Jonge et al., 2012) caused no significant stunting on Aibou plants, confirming that this remains a race 2 strain, while strains DVD161 and DVD3 caused significant stunting, implying that these strains should actually be assigned to race 3. As expected, the race 1 strain JR2 did not cause stunting on Aibou plants, which can at least partially be attributed to VdAve1 effector recognition by the *Ve1* gene product in these plants. However, the finding that a transgenic *VdAve1* deletion line (JR2ΔAve1; de Jonge et al., 2012) caused significant stunting on *Ve1*-transgenic Moneymaker and not on Aibou plants, suggests that the JR2 strain encodes Av2. However, this interpretation needs to be made with caution given that the virulence of the *VdAve1* deletion strain on tomato is severely compromised (de Jonge et al., 2012), as can be observed on Moneymaker plants in our assays (Fig. 1b). This observation, combined with the observation that stunting on Aibou plants by any race 3 strain is generally less than stunting on Moneymaker plants (Fig 1b, d), could indicate that basal defence against Verticillium wilt is enhanced in Aibou plants, and thus that incompatibility of the *VdAve1* deletion strain may be due to enhanced basal defence rather than due to V2-mediated recognition of the JR2 strain.

### Comparative genomics identifies *Verticillium dahliae Av2* candidates

Based on our phenotyping results, showing that unambiguous race assignment was not possible for every strain, four different groupings into race 2 and 3 strains with different degrees of stringency were made (Table 2). In the most stringent approach (I), comparative genomics was performed based on the six strains with the clearest phenotyping results, i.e. the race 2 strains TO22, UD-1-4-1 and GF-CA2 and the race 3 strains HOMCF, GF-CB5 and GF1192 (Usami et al., 2017). In a second approach (II), these strains were supplemented with the race 2 strain DVDS26 and race 3 strains DVD161 and DVD3. In a third approach (III), strains of approach II were supplemented with the previously assigned race 2 strain GF1207 and the previously assigned race 3 strain VT-2A (Usami et al., 2017), for which the phenotyping was not unambiguous in our assays. In the last approach (IV) we used the JR2 genome assembly (Faino et al., 2015) as a reference for comparative genomics with the selection of race 2 and race 3 strains as defined in approach II.

Besides assembly of strain JR2 (Faino et al., 2015), genome assemblies were also available for strains DVDS26, DVD161 and DVD3, albeit that these assemblies were highly fragmented as these were based on Illumina short-read sequencing data (de Jonge et al., 2012) (Table 1). In this study, we determined the genome sequences of four additional *V. dahliae* strains that belong to race 2, namely TO22, UD1-4-1, GF1207 and GFCA2, and four additional race 3 strains, namely GFCB5, GF1192, VT2A and HOMCF, with Oxford Nanopore sequencing Technology (ONT) using a MinION device (Table 1). For each strain, ∼2-3 Gb of sequence data was produced, representing 50-100x genome coverage based on the ∼35 Mb gapless reference genome of *V. dahliae* strain JR2 (Faino et al., 2015). Subsequently, we performed self-correction of the reads, read trimming and genome assembly, leading to genome assemblies ranging from 18 contigs for strain UD1-4-1 to 69 for strain GF1207 (Table 1).

To perform comparative genomics, for each of the approaches self-corrected reads from the selected *V. dahliae* race 3 strains were mapped against the assembly of *V. dahliae* strain TO22 (approaches I-III) or the telomere-to-telomere assembly of strain JR2 (approach IV) (Li, 2013) and regions that were not covered by race 3 reads were retained (Table 2). Next, self-corrected reads from the selected race 2 strains were mapped against the retained reference genome-specific regions that are absent from the race 3 strains. Sequences that were found in every race 2 strain were retained as candidate regions to encode the Avr molecule. Sequences that are shared by the *V. dahliae* strain TO22 reference assembly and the race 2 strains, and that are absent from the race 3 strains, were mapped against the *V. dahliae* strain JR2 genome assembly, and common genes were extracted. Sequences that did not map to the *V. dahliae* strain JR2 genome assembly were *de novo* annotated and signal peptides for secretion at the N-termini of the encoded proteins were predicted to identify potential effector genes.

Our strategy identified 670 kb of race 2-specific regions, containing 122 genes of which eight encode putative secreted proteins, for approach I (Table 2). For approach II, the addition of three strains reduced the target regions to 660 kb, containing 115 genes, of which six encode secreted proteins (Table 2). For approach III, the further addition of two strains for which the phenotyping was somewhat ambiguous slightly reduced the amount of candidate regions to 563 kb comprising 110 genes. However, the same six putatively secreted proteins remained as candidates, and the list of Avr candidates effectively remained the same as for approach II (Table 2). The last comparison in approach IV, based on the reference assembly of the *V. dahliae* JR2 strain, identified 277 kb of sequence as race 2-specific with only two genes that are predicted to encode secreted proteins; *XLOC_00170* (*VDAG_JR2_Chr4g03680a*) and *evm*.*model*.*contig1569*.*344* (*VDAG_JR2_Chr4g03650a*, further referred to as *Evm_344*), both of which were previously recognized as being among the most highly expressed effector genes during colonization of *Nicotiana benthamiana* plants (de Jonge et al. 2013; Faino et al., 2015).

### *In planta* expression of *Av2* candidates

We anticipate that the genuine *Av2* gene should not necessarily be expressed in *N. benthamiana* (de Jonge et al. 2013), but particularly in tomato. Real-time PCR analysis on a time course of tomato plants inoculated with the *V. dahliae* JR2 strain revealed that the two candidate genes are expressed during tomato colonization, with a peak in expression around 7 days post inoculation, whereas little to no expression could be recorded upon growth *in vitro* (Fig. 2a). Both genes are similarly expressed in *V. dahliae* strain TO22, albeit that the expression peaks slightly later, at 11 dpi (Fig. 2b). However, whereas the expression level of both genes is similar in *V. dahliae* strain JR2, *Evm_344* is higher expressed than *XLOC_00170* in *V. dahliae* strain TO22. Importantly, none of the six additional effector gene candidates that we identified in our comparative genomics approaches I-III and that are absent from *V. dahliae* strain JR2 are expressed *in planta* in *V. dahliae* strain TO22 (Fig. 2b). Thus, based on the transcriptional profiling these six effector genes can be disqualified as *Av2* candidates, and only two genes that display an expression profile that can be expected for a potential avirulence effector gene remain; *XLOC_00170* and *Evm_344*.

**Figure 2.**
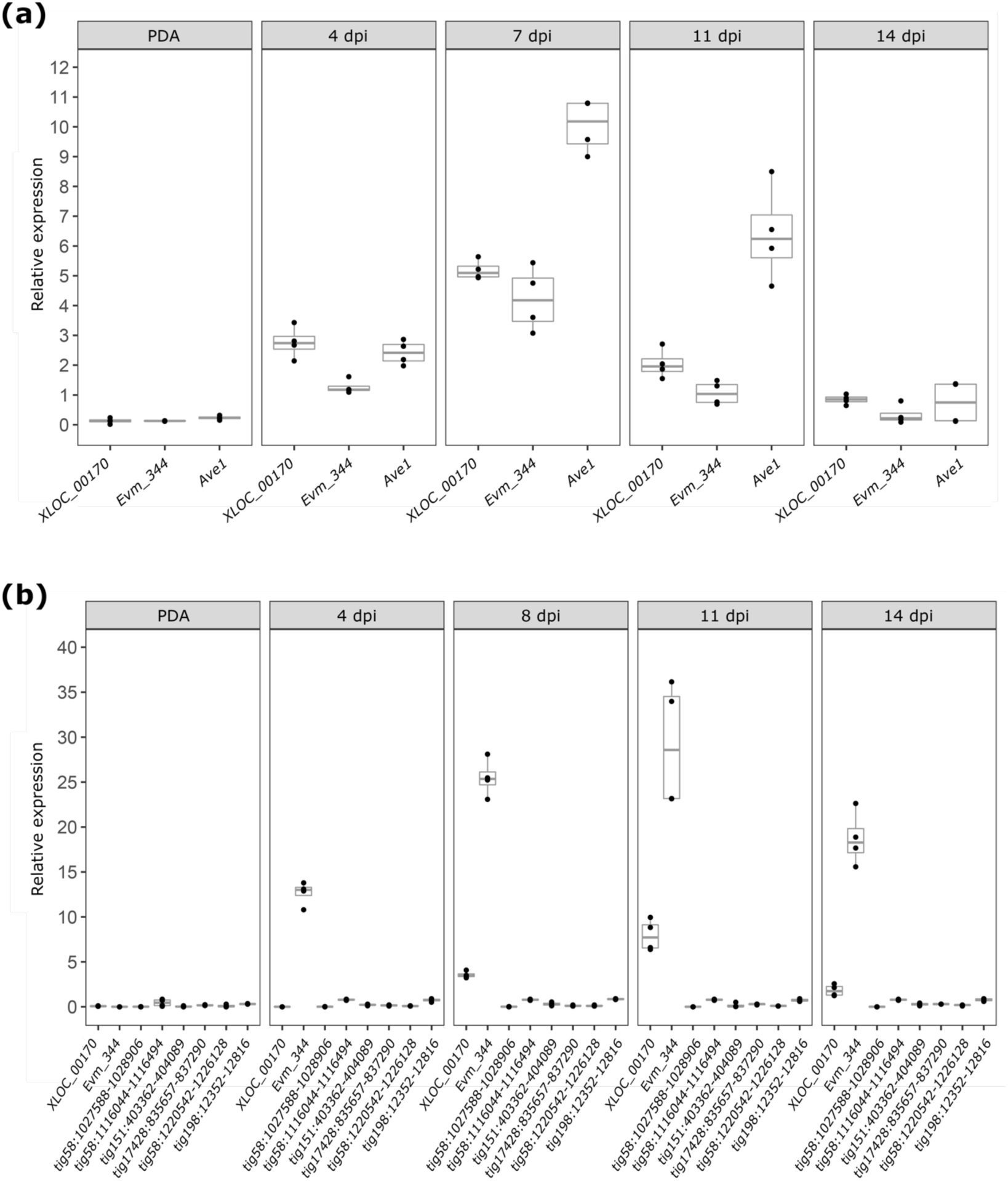
Expression of *V. dahliae* candidate avirulence effector genes *in vitro* and during colonization of tomato plants. To assess *in planta* expression, twelve-day-old tomato cv. Moneymaker seedlings were root-inoculated with *V. dahliae* strain JR2 **(a)** or strain TO22 **(b)**, and plants were harvested from 4 to 14 days post inoculation (dpi), while conidiospores were harvested from five-day-old cultures of *V. dahliae* on potato dextrose agar (PDA) to monitor *in vitro* expression. Real-time PCR was performed to determine the relative expression of *XLOC_00170, Evm_344* and the race 1-specific effector gene VdAve1 as a positive control (de Jonge et al., 2012) for strain JR2, using *V. dahliae GAPDH* as reference **(a)**. Similarly, the relative expression of *XLOC_00170, Evm_344* and six additional avirulence effector genes for strain TO22, using *V. dahliae GAPDH* as reference **(b)**.

### Occurrence of *Av2* candidates in a *V. dahliae* population

To determine which of the two remaining effector genes encodes Av2, a fungal isolate that carries either of the candidates separately should be tested for the ability to cause disease on Aibou plants that carry *V2*. Therefore, we attempted to identify *V. dahliae* isolates that carry only one of the two effector candidates. To this end, presence-absence variations (PAV) were assessed in a collection of 52 previously sequenced *V. dahliae* strains (Fig. 3; de Jonge et al., 2012; Faino et al., 2015; Fan et al., 2018; Gibriel et al., 2019). Remarkably, both effector genes always co-occurred in 17 of the isolates, including the four race 2 isolates that were sequenced in this study, and we were not able to identify any isolate that carries only a single of the two effector genes (Fig. 3).

**Figure 3.**
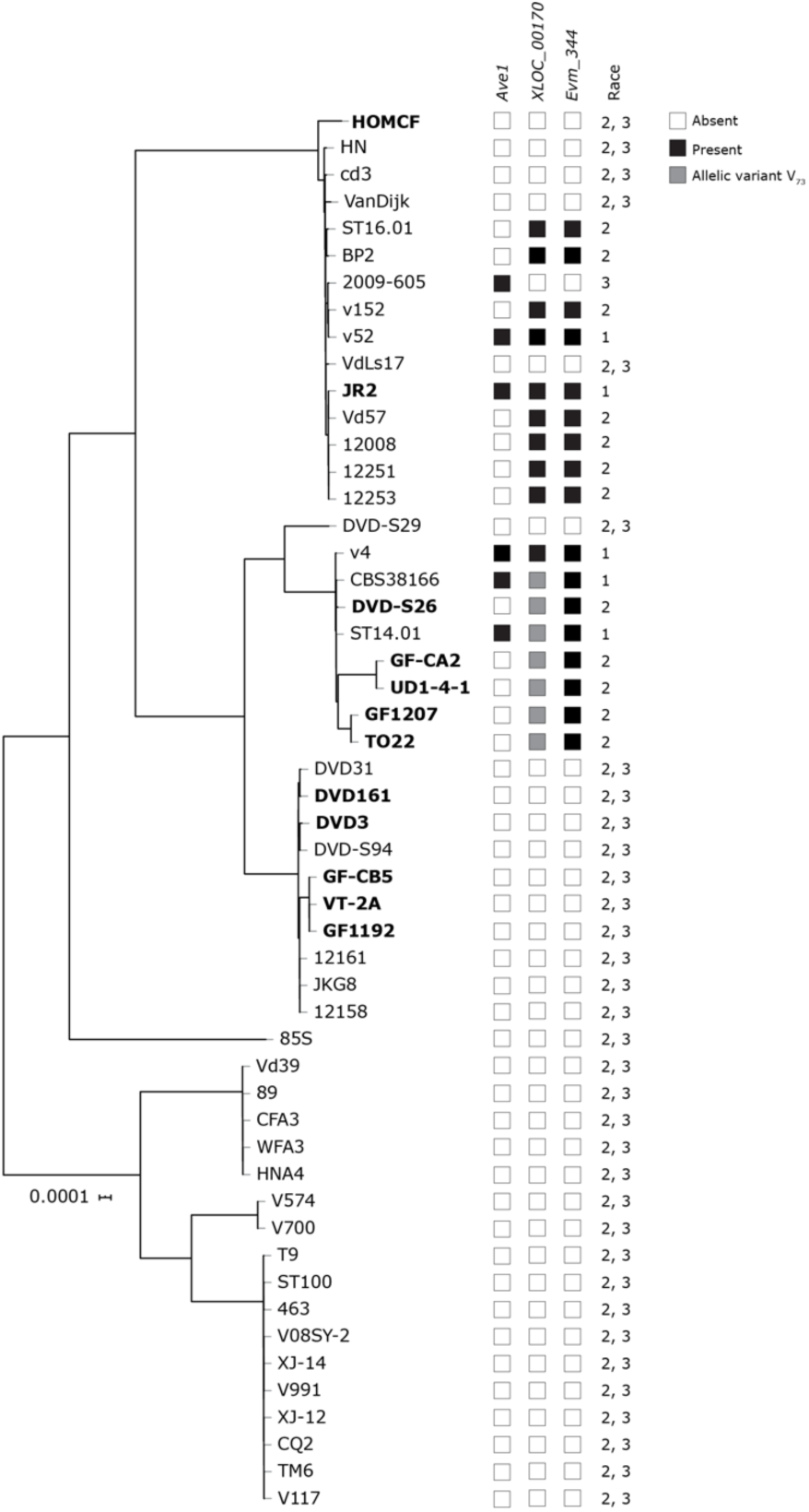
Phylogenetic tree of sequenced *V. dahliae* strains with indication of presence-absence variation in race 1 and race 2 effector candidates. Strains that were phenotyped and included in any of the comparative genomics approaches (Table 2) are shown in bold. Presence of the avirulence gene VdAve1, avirulence candidate genes *XLOC_00170* and *Evm_344*, and the race designation based on presence or absence of these genes are indicated. Phylogenetic relationships between sequenced *V. dahliae* strains were inferred using Realphy (Langmead & Salzberg, 2012).

To assess the phylogenetic relationships between strains that carry the two *Avr* candidates and strains lacking these candidates a phylogenetic tree was generated, showing that the strains can be grouped into three major clades, two of which carry strains that contain the two *Avr* candidate genes. However, within these clades closely related strains occur that lost the effector genes, suggesting the occurrence of multiple independent losses (Fig. 3). However, overall, no obvious phylogenetic structure is apparent with respect to effector presence within the *V. dahliae* population.

The co-occurrence of both *Avr* effector candidate genes suggests that they are physically linked in the genome. Therefore, the genomic organization surrounding these two effector gene candidates was investigated based on the gapless genome assembly of *V. dahliae* strain JR2 (Faino et al., 2015). Indeed, both genes locate in close proximity of each other, separated by only two additional genes, in a lineage-specific (LS) region on chromosome 4 (Fig. 4). Furthermore, as typically observed in LS regions that are enriched in repetitive elements (de Jonge et al., 2013; Faino et al., 2016), the genes are surrounded by repetitive elements such as transposons that mostly belong to the class II long terminal repeat (LTR) retrotransposons (Fig. 4, Fig. 5). Typically, LS regions are characterized by the high abundance of PAV. As expected, the flanking genomic region (100 kb) are highly variable between *V. dahliae* strains (Fig. 5). Even though these two effector gene candidates are located in a plastic region, their close proximity suggests that it is unlikely to find naturally occurring *V. dahliae* isolates that carry only one of the two effector gene candidates, and consequently other strategies than testing natural isolates on Aibou plants for confirming which of the two candidates encodes Av2 should be pursued.

**Figure 4.**
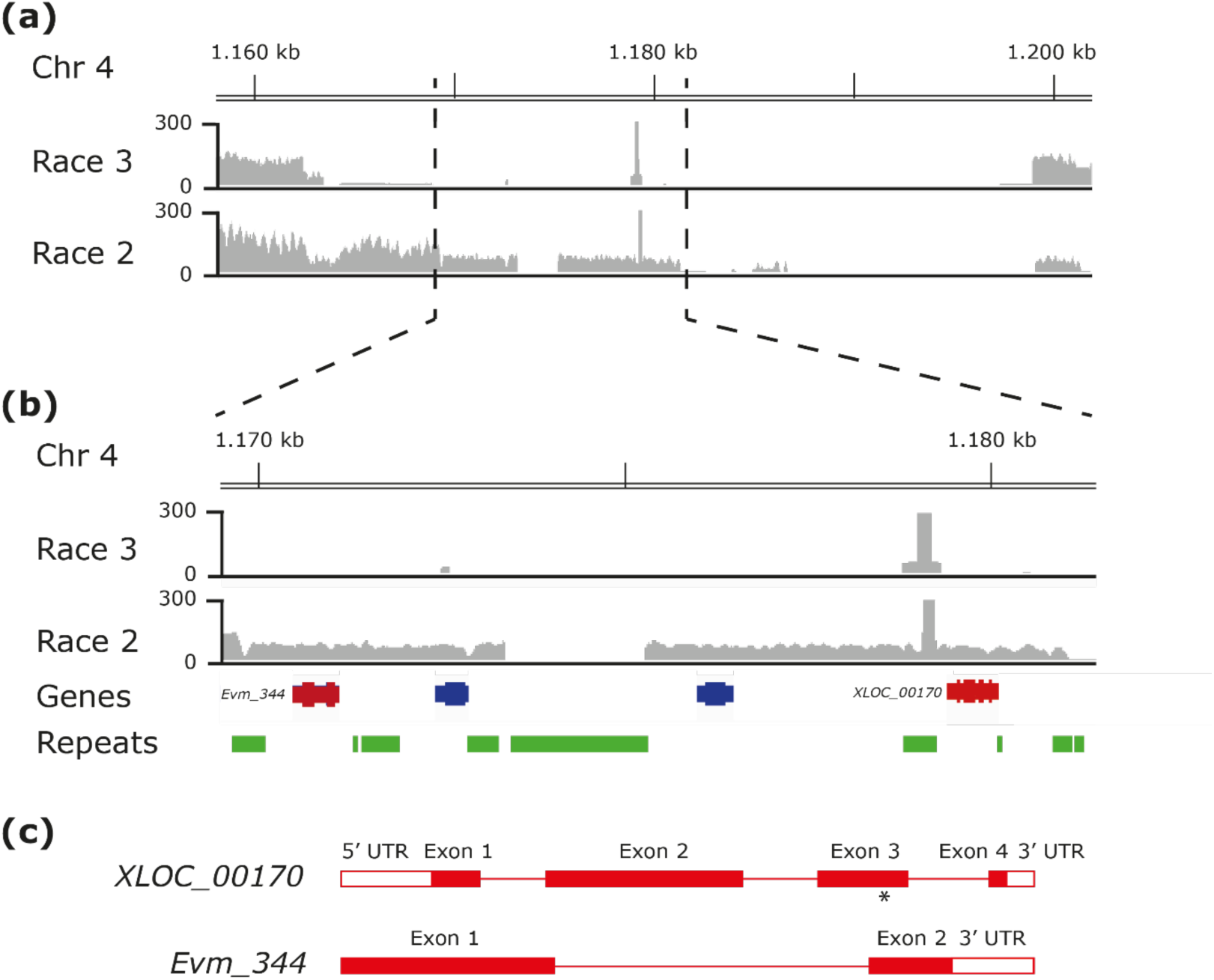
Genomic organization of the locus comprising the two candidate Avr effector genes in *V. dahliae* strain JR2. **(a)** Schematic representation of the genomic region on chromosome 4 that is largely lacking in the race 3 strains that are included in comparative genomics approach IV (Table 2). Reads mapping to the region are shown as coverage plots for race 3 and race 2 strains. **(b)** Close-up of the region surrounding the *Avr* effector gene candidates *XLOC_00170* and *Evm_344* that are indicated as red gene models. Other gene models are show in blue and repetitive elements in green. **(c)** Gene models for *XLOC_00170* and *Evm_344*. The asterisk indicates the approximate position of the single (A to T) nucleotide substitution in the *XLOC_00170* gene of 7 of the 17 isolates that carry the gene, leading to a single amino acid substitution (E_73_V).

**Figure 5.**
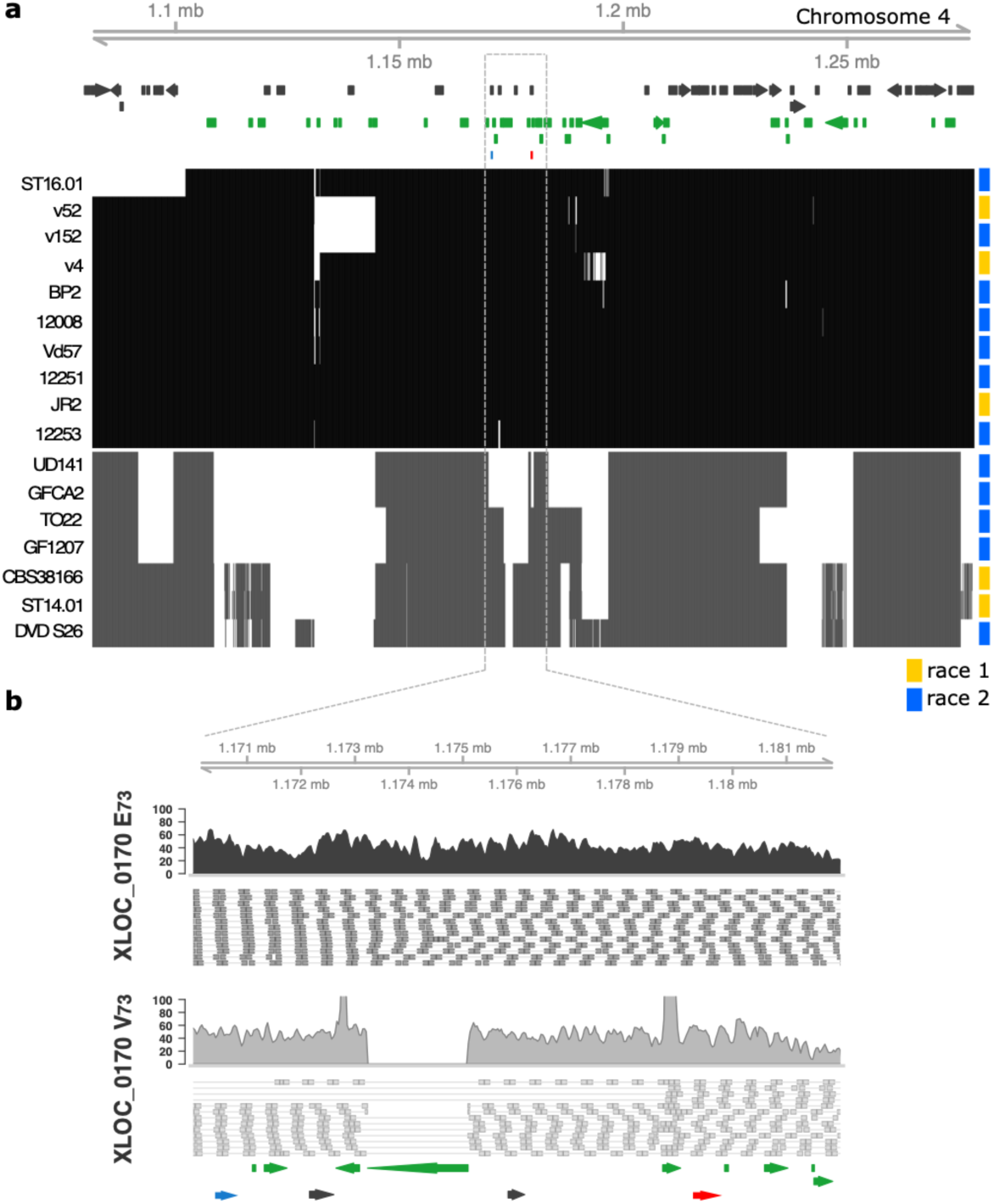
Presence-absence variation in the region surrounding the two candidate *Av2* genes. **(a)** Genomic region flanking the *Av2* candidate genes in 17 isolates detailed in **Figure 3**. The matrix shows the presence (black/grey) and absence (white) in 100 bp non-overlapping windows for XLOC_0170 variant E_73_ (black) and XLOC_0170 variant V_73_ grey. On top, annotated genes are displayed in black and repetitive elements in green, while *XLOC_00170* is displayed in red and *Evm_344* in blue. **(b)** Read coverage for *V. dahliae* strain JR2 that produces XLOC_0170 variant E_73_ and strain DVD-S26 that produces XLOC_0170 variant V_73_ depicting a transposable element deletion in isolates that produce the V_73_ variant.

### Expression in *V. dahliae* race 3 reveals that *XLOC_00170* encodes Av2

To identify which of the two candidates encodes Av2, a genetic complementation approach was pursued in which the two candidate genes were introduced individually into the *V. dahliae* race 3 strains GF-CB5 and HOMCF. Subsequently, inoculations were performed on a differential set of tomato genotypes, comprising Moneymaker plants, *Ve1*-transgenic Moneymaker plants (Fradin et al., 2009), and Aibou plants (Usami et al., 2017). As expected, the non-transformed race 3 strains GF-CB5 and HOMCF as well as the complementation lines containing *XLOC_00170* or *Evm_344* caused clear stunting of the universally susceptible Moneymaker as well as of the *Ve1-*transgenic Moneymaker plants (Fig. 6a, b). Interestingly, whereas non-transformed race 3 strains GF-CB5 and HOMCF as well as the complementation lines containing *Evm_344* caused clear stunting on Aibou plants, complementation lines expressing the *Avr* candidate *XLOC_00170* did not induce disease symptoms and could not induce stunting on these plants (Fig. 6a, b). As such, these complementation transformants of the race 3 strains GF-CB5 and HOMCF behaved essentially as the race 2 strain TO22 (Fig. 6a, b). Thus, these findings suggest that *XLOC_00170* encodes Av2. All visual observations of stunting were supported by quantifications of fungal biomass by real-time PCR (Fig. 6c). These measurements revealed that fungal biomass levels were only reduced on Aibou plants when inoculated with the race 2 strain TO22, and with the race 3 strains GF-CB5 and HOMCF that were complemented with *XLOC_00170*. Thus, our data collectively confirm that the reduced symptomatology is accompanied by significantly reduced fungal colonization and demonstrate that *XLOC_00170* encodes the race 2-specific avirulence effector Av2.

**Figure 6.**
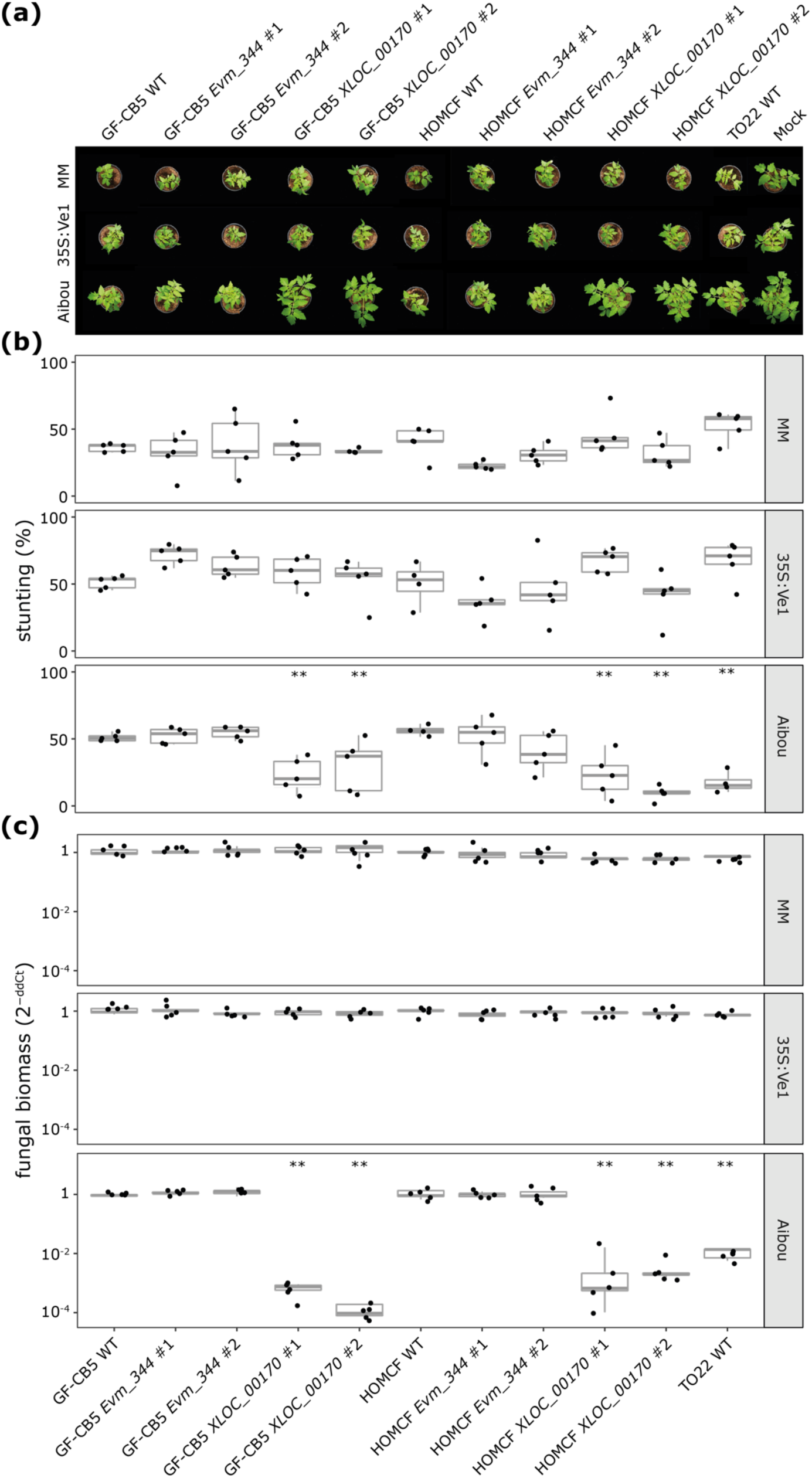
*XLOC_00170* encodes the avirulence effector Av2 that is recognized in *V2* plants. **(a)** Top pictures of Moneymaker plants that lack known *V. dahliae* resistance genes (MM), *Ve1*-transgenic Moneymaker plants that are resistant against race 1 and not against race 2 strains of *V. dahliae* (MM 35S:Ve1), and Aibou plants that carry *Ve1* and *V2* and are therefore resistant against race 1 as well as race 2 strains of the pathogen (Usami et al., 2017) inoculated with the race 3 WT strains GF-CB5 and HOMCF, and two independent genetic complementation lines that express *XLOC_00170* or *Evm_344*, and the race 2 strain TO22. **(b)** Quantification of stunting caused by the various *V. dahliae* genotypes on the various tomato genotypes as detailed for panel **(a)**. Each combination is represented by the measurement of five plants. **(c)** Quantification of fungal biomass with real-time PCR determined for the various *V. dahliae* genotypes on the various tomato genotypes as detailed for panel **(a)**. Each combination is represented by the fungal biomass quantification in five plants.

### *Av2* allelic variation

As many Avr effectors are under selection pressure, and thus often display enhanced allelic variation (Stergiopoulos et al., 2007), we assessed the 17 effector gene alleles that were identified in the collection of 52 previously sequenced *V. dahliae* strains (Fig. 3) for allelic variation. Whereas no allelic variants were found for *Evm_344*, two single allelic variants were found for *Av2*, concerning a single nucleotide polymorphism in exon 3 leading to a polymorphic amino acid at position 73. Whereas ten isolates carry a glutamic acid at this position (E_73_), seven other carry a valine (V_73_) (Fig. 7). Interestingly, strains carrying V_73_ are clustered in the same branch, suggesting that a single event caused the polymorphism (Fig. 3). Intriguingly, we noticed that all isolates carrying E_73_ carry an extra transposable element of the DNA/Tc-1 Mariner class in the upstream region of the *Av2* gene (Fig. 5). However, as strains GF-CA2, TO22, UD-1-4-1 DVDS26 and GF1207, that encode the Av2 variant with V_73_, as well as the VdAve1 deletion strain of JR2, that encode the variant with E_73_, are contained on Aibou plants, it needs to be concluded that both allelic variants are recognized by V2.

**Figure 7.**
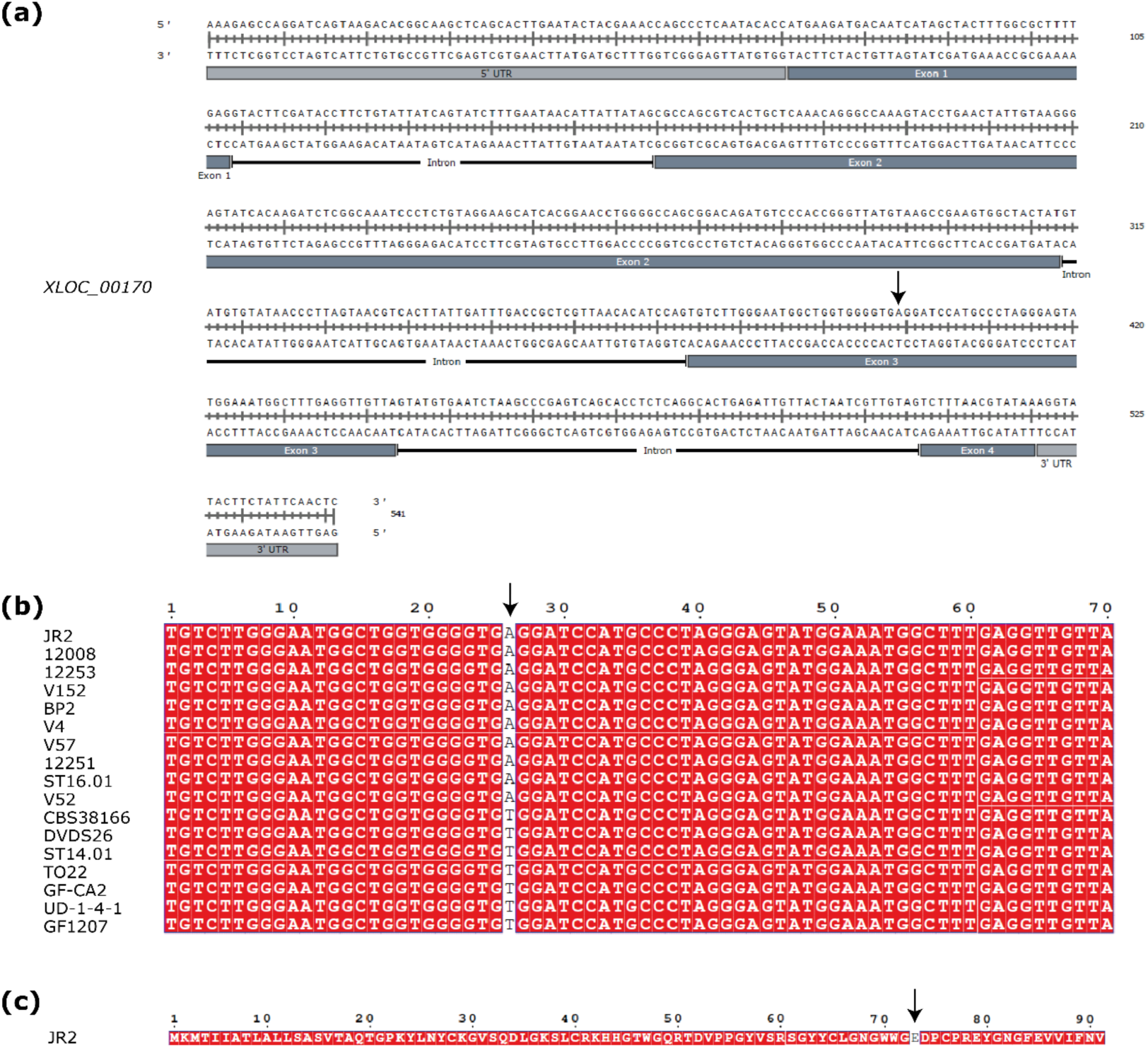
Allelic variation of the effector candidate gene *XLOC_00170* in sequenced strains carrying the effector. **(a)** Genomic sequence of effector candidate gene *XLOC_00170*, the arrow shows the position of the single nucleotide substitution found in particular strains. **(b)** Alignment of exon 3 of the effector candidate gene in the 17 strains containing the effector. The arrow shows the single nucleotide substitution that occurs in seven of the strains when compared with strain JR2. **(c)** Amino acid sequence of the effector candidate gene *XLOC_00170* as encoded by *V. dahliae* strain JR2 with E_73_ that is substituted by V in seven isolates indicated by an arrow.

## DISCUSSION

Historically, the identification of avirulence genes has been challenging for fungi that reproduce asexually, as genetic mapping cannot be utilized. However, since the advent of affordable genome sequencing, cumbersome and laborious methods to identify avirulence genes, that include functional screenings of fungal cDNAs or protein fractions for the induction of immune responses in plants, have been supplemented with comparative genomics and transcriptomics strategies (Gibriel et al., 2016). Less than a decade ago we have been the first to identify the first avirulence gene of *V. dahliae*, known as VdAve1 for mediating <u>a</u>virulence on *<u>Ve1</u>* plants, through a comparative population genomics strategy combined with transcriptomics by utilizing race 1 strains that were contained by the *Ve1* resistance gene of tomato, and resistance-breaking race 2 strains (de Jonge et al., 2013). In this study, we used a similar approach based on comparative population genomics of race 1 and 2 strains with race 3 strains to successfully identify XLOC_00170 as the Av2 effector that mediates avirulence on V2 plants. Intriguingly, besides VdAve1, XLOC_00170 has been identified previously as one of the most highly induced genes of *V. dahliae* during host colonization (de Jonge et al., 2013).

*Ve1* and the *V2* locus are the only two major resistance sources that have thus far been described in tomato against *V. dahliae* (Fradin et al., 2009; Usami et al., 2017). Since its initial introduction from a wild Peruvian tomato accession into cultivars in the 1950s (Deseret News and Telegram, 1955), *Ve1* has been widely exploited as it is incorporated in virtually every tomato cultivar today. Even though soon after the introduction of these cultivars resistance-breaking race 2 strains emerged, first in the USA (Alexander 1962; Robinson 1957), and soon thereafter also in Europe (Cirulli, 1969; Pegg & Dixon, 1969), *Ve1* is still considered useful for Verticillium wilt control today. An important factor that contributes to the durability of resistance is the fitness penalty for the pathogen upon losing the corresponding avirulence factor (Brown, 2015). Indeed, the VdAve1 effector contributes considerably to *V. dahliae* virulence on tomato, which explains why race 2 strains that lack *VdAve1* are generally less aggressive (de Jonge et al., 2012). Based on our current observations, differences in aggressiveness between race 2 and race 3 strains on Moneymaker plants are not obvious (Fig. 1), and also genetic complementation of race 3 strains with *XLOC_00170* did not lead to a striking increase in aggressiveness on Moneymaker plants (Fig. 6), suggesting that the contribution of *Av2* to *V. dahliae* virulence under the conditions tested in this study is modest at most.

Thus far, V2 resistance has been exploited scarcely when compared with Ve1, as it has only been introduced in a number of Japanese rootstock cultivars since 2006 (Usami et al., 2017). Previously, *V2* resistance-breaking race 3 strains have been found in several Japanese prefectures on two separate islands (Usami et al., 2017). Intriguingly, our genome analyses demonstrate that race 3 strains that lack *Av2* are ubiquitous and found worldwide, as our collection of sequenced strains comprises specimens that were originally isolated in Europe, China, Canada, and the USA. Arguably, most of these race 3 strains arose in the absence of *V2* selection by tomato cultivation. It is conceivable that, similar to Ve1 homologs that are found in other plant species besides tomato (Song et al., 2017), functional homologs of *V2* occur in other plant species as well, which may have selected against the presence of *Av2* in many *V. dahliae* strains. However, as long as the identity of *V2* remains enigmatic this hypothesis cannot be tested.

Like *VdAve1*, also *Av2* resides in an LS region of the *V. dahliae* genome, albeit in another region on another chromosome. Typically, these LS regions are gene-sparse and enriched in repetitive elements, such as transposons, causing these regions to be highly plastic which is thought to mediate accelerated evolution of effector catalogues (de Jonge et al., 2013; Faino et al., 2016; Cook, et al., 2020). We previously demonstrated that *VdAve1* has been lost from the *V. dahliae* population multiple times, and to date only PAV has been identified as mechanism to escape *Ve1*-mediated immunity (de Jonge et al., 2013, 2012; Faino et al., 2016). Similarly, our phylogenetic analysis reveals that *Av2* has been lost multiple times independently, and although we identified two allelic variants, both variants are recognized by V2. Consequently, PAV remains the only mechanism to overcome *V2*-mediated immunity thus far. Despite the observation that PAV is the only observed mechanism for *V. dahliae* to overcome host immunity, pathogens typically exploit a wide variety of mechanisms, ranging from single nucleotide polymorphisms (Joosten et al., 1994) to altered expression of the avirulence gene (Na and Gijzen, 2016). Nevertheless, avirulence gene deletion to overcome host immunity is common and has been reported for various fungi, including *C. fulvum* (Stergiopoulos et al., 2007), *Fusarium oxysporum* (Schmidt et al., 2016), *Leptosphaeria maculans* (Gout et al., 2007; Parlange et al., 2009), and *Magnaporthe oryzae* (Pallaghy et al., 1994; Zhou et al., 2007).

It was previously demonstrated that frequencies of single nucleotide polymorphisms (SNPs) are significantly reduced in the area surrounding the VdAve1 locus when compared with the surrounding genomic regions (Faino et al., 2016), which was thought to point towards recent acquisition through horizontal transfer (de Jonge et al., 2012). However, we recently noted that enhanced sequence conservation through reduced nucleotide substitution is a general feature of LS regions in *V. dahliae* (Depotter et al., 2019). Although a mechanistic underpinning is still lacking, we hypothesized that differences in chromatin organization may perhaps explain this phenomenon. Interestingly, while DNA methylation is generally low and only present at TEs, only TEs in the core genome are methylated while LS TEs are largely devoid of methylation (Cook et al., 2020). Furthermore, TEs within LS regions are more transcriptionally active and display increased DNA accessibility, representing a unique chromatin profile that, likely, contributes to the plasticity of these regions (Cook et al., 2020; Faino et al., 2016). Possibly, the increased DNA accessibility makes that genes that reside in these regions tend to be overrepresented during expression *in planta*, and *VdAve1* as well as *Av2* belong to the most highly expressed genes during host colonization (de Jonge et al., 2013). Despite the repressed SNP frequencies in LS regions, we found two allelic variants in the *V. dahliae* population. Intriguingly, the isolates that carry the Av2 variant E_73_ carry a nearby insertion of DNA/Tc-1 Mariner transposon. However, a mechanistic link between the transposon insertion and the occurrence of the allelic variant is not obvious.

Our identification of *Av2* concerns the cloning of only the second avirulence gene of *V. dahliae*. This identification permits its use as a functional tool for genetic mapping of the *V2* gene. Typically, *V. dahliae* symptoms on tomato display considerable variability, and disease phenotyping is laborious. Possibly, injections of heterologously produced Av2 protein can be used to screen tomato plants in genetic mapping analyses, provided that such injections result in a visible phenotype such as a hypersensitive response. Similar effector-assisted resistance breeding has previously been used successfully identify resistance sources in tomato against the leaf mould pathogen *Cladosporium fulvum* (Lauge et al., 1998; Takken et al., 1999) and potato against the late blight pathogen *Phytophthora infestans* (Du et al., 2015; Vleeshouwers and Oliver, 2014). The identification of *Av2* can furthermore be exploited for race diagnostics of *V. dahliae* to determine whether cultivation of resistant tomato genotypes is useful, but also to monitor *V. dahliae* population dynamics and race structures. Arguably, based on the identification of avirulence genes, rapid in-field diagnostics can be developed to aid growers to cultivate disease-free crops.

## ACKNOWLEDGEMENTS

EACC and DET acknowledge receipt of PhD fellowships from CONACyT. Work in the laboratories of BPHJT and MFS is supported by the Research Council for Earth and Life Science (ALW) of the Netherlands Organization for Scientific Research (NWO). BPHJT acknowledges support by the Deutsche Forschungsgemeinschaft (DFG, German Research Foundation) under Germany’s Excellence Strategy – EXC 2048/1 – Project ID: 390686111. Part of the work was funded by Foundation Topconsortium voor Kennis en Innovatie (TKI) Starting Materials, project number 1409-026.

## AUTHOR CONTRIBUTIONS

JPV, TU, MFS and BPHJT conceived the study; EACC, JPV, DET, MFS and BPHJT designed experiments; EACC, JPV, DET performed experiments; EACC, JPV, DET, HJS, YB, MFS and BPHJT analyzed data, EACC, JPV and BPHJT wrote the manuscript; MFS and BPHJT supervised the project, all authors discussed the results and contributed to the final manuscript.

